# Fibroblasts sense spatial proximity via an EGFR–CREB5 axis to restore quiescent synovial lining in remission rheumatoid arthritis

**DOI:** 10.64898/2025.12.10.693501

**Authors:** Sonia Presti, Ksenia S. Anufrieva, Ce Gao, Sean A Prell, Kartik Bhamidipati, Vikram Khedgikar, Philip E Blazar, Jeffrey K Lange, Morgan H Jones, Michael B Brenner, Ellen M Gravallese, Ilya Korsunsky, Mihir D Wechalekar, Shideh Kazerounian, Kevin Wei

**Author notes:** Corresponding authors Contact: Kevin Wei, MD, PhD Address: 60 Fenwood Rd., Hale BTM 6016cc, Boston, MA 02115, USA. These authors contributed equally (Sonia Presti, Ksenia S. Anufrieva, Ce Gao).

## Abstract

Rheumatoid arthritis (RA) is a chronic inflammatory disease where the synovial lining membrane undergoes pathological changes resulting in joint destruction. In healthy joints, the synovial lining is essential for joint homeostasis, forming a selective barrier and secreting lubricating molecules, yet the mechanisms that restores homeostatic synovial lining during RA remission remains poorly understood. Here, we applied spatial transcriptomics to examine biopsies of RA patients in remission to identify a mechanism that orchestrates a phenotypic switch specifying synovial quiescent lining fibroblast differentiation. Spatial transcriptomics revealed a proximity-sensing program where at low cell-density, fibroblasts adopt proliferative and fibrotic transcriptional state characterized by expression of *MKI67*, *COL1A2* and *COL6A2*, whereas at high cell-density, fibroblasts induce a quiescent lining fibroblast transcriptional program characterized by *PRG4*, *CLU* and *PDPN*. Mechanistically, fibroblasts sense spatial proximity through HB-EGF–EGFR signaling, which leads to phosphorylation of transcription factor CREB5. Perturbation of the EGFR-CREB5 axis abolishes fibroblast proximity-sensing and blocks synovial lining fibroblast differentiation. Conversely, EGFR activation by the ligand HB-EGF or pharmacologic activation of CREB5 is sufficient to induce synovial lining fibroblast differentiation. Together, our findings define a novel spatial proximity-sensing pathway underlying a return to homoeostatic fibroblast function during RA remission. By sensing their spatial proximity to neighboring fibroblasts, synovial fibroblasts translate these positional cues into signals that lead to restoration of normal, steady-state synovial lining membrane.

## Introduction

Rheumatoid arthritis (RA) is a chronic autoimmune disease that causes persistent inflammation of the synovial membrane, leading to cartilage damage, bone erosion, and joint deformity^1,2^. The synovium is a specialized connective tissue that protects the joint and maintains lubrication. In healthy joints, synovial lining fibroblasts and macrophages form a protective barrier and promote synovial membrane homeostasis^3,4^. In RA, the synovium undergoes hyperplasia and becomes invasive, forming a pannus that erodes cartilage and bone ^5^. Recent single-cell and spatial single-cell transcriptomic studies have shown that fibroblasts in the synovium are not uniform but form distinct populations with specific functions ^6–9^. Synovial lining fibroblasts sit at the surface of the synovium and express genes crucial for barrier and lubrication functions, including *PRG4* and *CLU*, while sublining fibroblasts lie deeper in the synovial tissue and express inflammatory and matrix-remodeling genes including *THY1*, *CXCL12*, and *COL1A2* ^10,11^. In active RA, synovial lining fibroblasts exhibit a pro-inflammatory phenotype ^9^ in response to inflammatory cytokines produced by infiltrating immune cells. Although most research has focused on fibroblasts during active RA, recent studies^12,13^ have begun to explore their role in remission, when inflammation resolves and the synovium regains a more normal structure. Analyses of synovial tissue from patients in drug-free remission show restoration of a thin, PRG4⁺ lining layer similar to that of healthy joints ^10,12,14^.

Anatomical location is an important driver of fibroblast heterogeneity ^15,8,16^. In the synovium, fibroblast positional memory is shaped by epigenics ^17,18^ and microenvironmental cues ^8,19^. Synovial fibroblasts rely on local environmental signals to establish and maintain their transcriptional state in synovial tissue, a fibroblast feature we termed as fibroblast positional identity ^8,16^. Synovial fibroblasts rapidly lose their positional identity when isolated from their native tissue microenvironment and de-differentiate *ex vivo* ^8^. We have previously demonstrated a crucial role for vascular endothelial-derived Notch signaling in driving sublining fibroblast expansion in RA as sublining fibroblasts sense the presence of neighboring endothelial cells through NOTCH3 to differentiate into vascular fibroblasts ^8,16^, a key pathological process underlying fibroblast expansion in active RA^8,10,20^. However, the microenvironment cues that specify the positional identity of steady-state synovial lining fibroblasts during RA remission remains unknown.

The rapid re-establishment of an organized synovial lining during RA remission suggests that, as inflammation resolves, synovial fibroblasts rely on spatial cues within their microenvironment to return to a quiescent transcriptional state. To define the molecular pathway that specifies and stabilizes quiescent lining fibroblasts, we combined high-resolution spatial transcriptomic profiling of synovial biopsies from RA patients in remission with controlled two- and three-dimensional systems that manipulate cell density and fibroblast proximity-sensing. These complementary approaches uncovered an unexpected requirement for spatial proximity in instructing fibroblast positional identity and promoting quiescence. Building on these observations, our findings identify a spatial HB-EGF–EGFR–CREB5 pathway that guides synovial fibroblasts in rebuilding the homeostatic lining during remission.

## Results

### Synovial fibroblasts exhibit cell density-associated gene expression

Our previous study demonstrated that synovial lining fibroblasts rapidly lose their transcriptional identity and transition toward a de-differentiated transcriptional state when removed from their native synovial tissue microenvironment ^8^. To identify the mechanisms underlying the restoration of quiescent synovial lining fibroblast state, we analyzed high-resolution spatial transcriptomic data from synovial biopsies of RA patients in clinical remission as defined by DAS28<2.6 after 6 months of treatment with either triple conventional synthetic disease-modifying anti-rheumatid drugs (csDMARD) therapy or adalimumab^19^. We obtained pre-annotated synovial fibroblasts (113,803 cells) from spatial transcriptomic data of synovial biopsies from 10 RA patients in remission. A comprehensive analysis identified two predominant fibroblast subtypes within the synovial tissue: *PRG4*+ lining fibroblasts and *THY1*+ sublining fibroblasts (**Fig. 1A** and **B**). Consistent with our prior studies^10^, lining fibroblasts and sublining fibroblasts exhibit distinct transcriptomic profiles, where sublining fibroblasts demonstrated an activated fibrogenic transcriptional program, marked by upregulation of transforming growth factor-beta (TGF-β) pathway components (**Fig. 1C**)^9^. In agreement with published single-cell transcriptomic atlas of RA synovia^10^, lining fibroblasts in remission expressed canonical markers *PRG4*, *PDPN*, and *CLU*, whereas sublining fibroblasts exhibited elevated expression of activated fibroblast marker *CDH11* and fibrogenic markers *COMP* and *COL5A1* (**Fig. 1D**).

**Figure 1.**
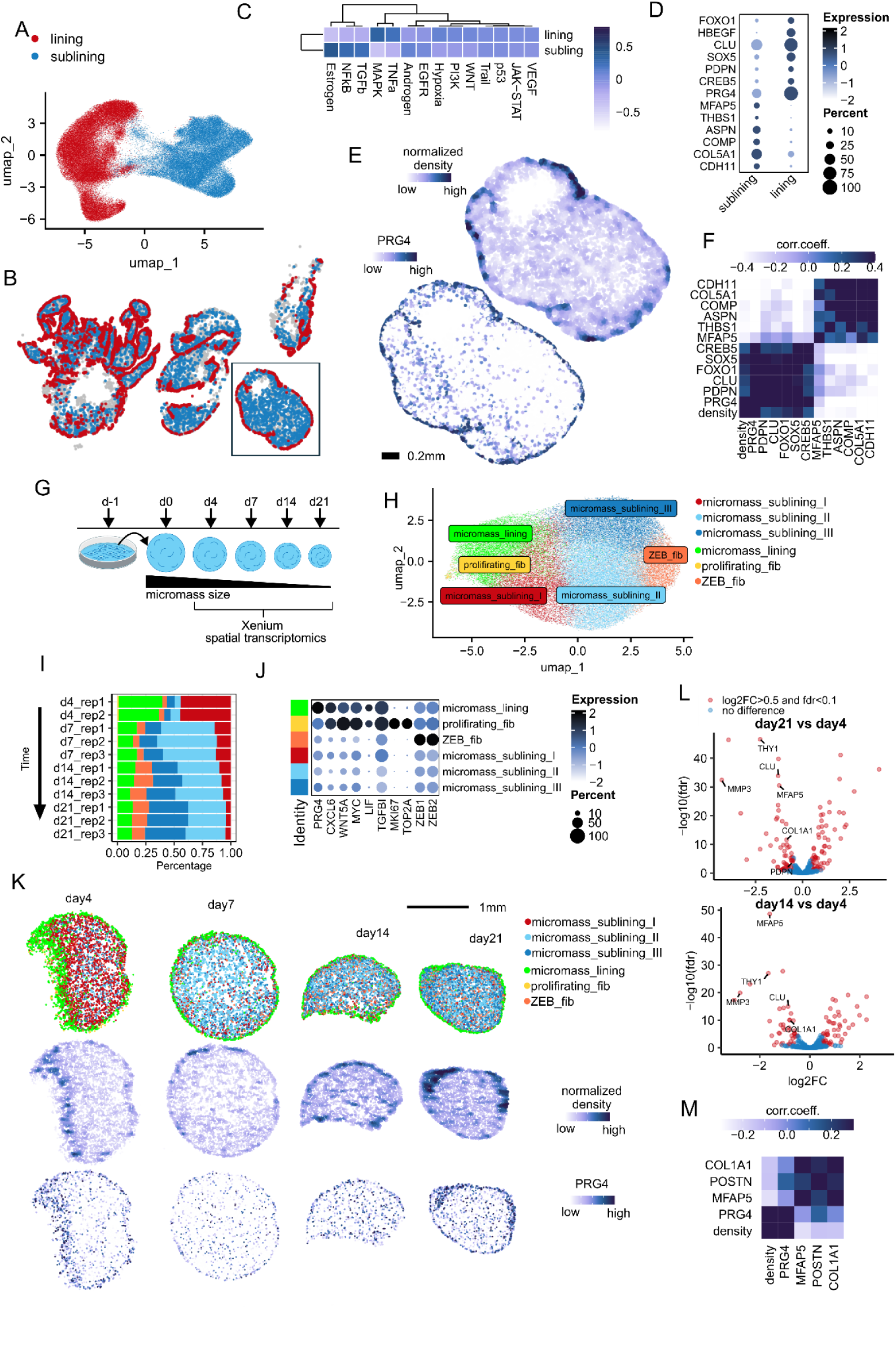
Spatial single cell profiling of synovial fibroblasts in RA tissue and micromass cultures **A.** UMAP visualization of synovial fibroblast subpopulations in RA tissue. **B.** Spatial distribution of lining fibroblasts (red), sublining fibroblasts (blue), and other cell types (grey) within synovial tissue sections from RA patients. **C.** PROGENy pathway analysis across synovial fibroblast subpopulations. **D.** Expression levels of marker genes in lining and sublining fibroblast populations. Dot size represents percent of cells expressing each gene, while color intensity indicates expression level. **E.** Cell density analysis of synovial tissue. The top image shows normalized fibroblast density distribution. The bottom image displays *PRG4* expression in the same section. **F.** Spatial correlation analysis between gene expression and density of fibroblasts in tissue. Heatmap showing correlation between fibroblast marker genes and normalized fibroblast cell density. **G.** Schematic representation of high-density micromass culture methodology **H.** UMAP visualization of fibroblast subclusters in micromass cultures. **I-J.** Temporal changes in fibroblast populations during micromass cultivation. (**I**) Stacked bar chart showing the proportion of each fibroblast subcluster across timepoints and replicates. (**J**) Dot plot showing marker gene expression across fibroblast subclusters identified during analysis of spatial single cell data. **K.** Spatial distribution of fibroblast subclusters in micromass cultures across cultivation timepoints. Top row shows cell identity (color-coded by cluster); middle row displays normalized cell density; bottom row shows *PRG4* expression. Fibroblast clusters are colored according to their classification as defined in Fig. 1H. **L.** Differential gene expression analysis of fibroblasts inside micromass cultures. Volcano plots comparing day 21 vs day 4 (top) and day 14 vs day 4 (bottom). **M.** Correlation heatmap showing relationships between cell density and marker gene expression in micromass systems.

Given that our dataset included both gene expression and spatial localization data for the synovial fibroblasts, we analyzed their spatial distribution pattern within the synovial tissue. To quantify the spatial organization, we computed fibroblast density using kernel density estimation with Diggle’s bandwidth selection and edge correction across the entire tissue section. This density map was then interpolated to assign a local fibroblast density value to each cell, regardless of cell type. Our analysis revealed that lining fibroblasts were predominantly localized in regions characterized by high local fibroblast density(**Fig. 1E**). Additionally, spatial correlation of gene expression revealed a positive association between canonical lining markers and fibroblast density, while non-canonical sublining genes were found to be enriched in areas of lower density (**Fig. 1F**).

To explore the relationship between cell density and fibroblast gene expression, we developed an alternative approach using synovial fibroblast micromass organoids. In this system, high-density micromasses are generated from dissociated synovial tissue fibroblasts that are expanded in culture and subsequently self-assemble into three-dimensional organoid structures^8,21^ (**Fig. 1G**). Over a 21-day period, we noticed that synovial organoids condense into smaller, compacted spheroids, forming a dense fibroblast-layer at the edges of the organoids (**Fig. 1K**). Spatial single-cell analysis revealed two fibroblast clusters localized at the edges of each micromass organoid: micromass_lining and proliferating_fib (**Fig. 1H** and **Suppl Table. 1**). The micromass_lining cluster expressed lining markers *PRG4* and *CXCL6* along with additional genes including *WNT5A*, *LIF*, *MYC*, and *TGFß1* (**Fig. 1J** and **Suppl Table. 1**). The proliferating_fib cluster shared this molecular signature but also expressed proliferation markers, including *MKI67* and *TOP2A* (**Fig. 1J**). While the micromass_lining cluster persisted throughout the entire culture period, the proliferating_fib cluster was observed only at the earliest time point (day 4) (**Fig. 1I**), suggesting that the micromass culture system supports maintenance of the lining fibroblast phenotype.

Spatial analysis also identified four distinct fibroblast clusters within the interior of the micromass: ZEB_fib, micromass_sublining_I, micromass_sublining_II, and micromass_ sublining_III (**Fig. 1H** and **I**). These clusters exhibited dynamic changes over time: sublining_I predominated at day 4, sublining_II at day 7, and sublining_III at days 14 and 21 (**Fig. 1I**). Due to the limited gene coverage of the spatial panel (477 genes), definitive markers distinguishing these subpopulations could not be identified (**Fig. 1J** and **Suppl Table. 1**). To assess temporal gene expression changes among internal fibroblast populations, we performed pseudo-bulk differential expression analysis by aggregating transcriptional data from all non-peripheral fibroblast clusters. This analysis revealed that classical myofibroblast/sublining markers such as *THY1*, *COL1A1*, and *MFAP* decreased over time, particularly at days 14 and 21 (**Fig. 1L** and **M**). This finding is consistent with our previous study demonstrating that endothelium-derived Notch signaling is required to maintain sublining fibroblast differentiation ^8^. To explore the underlying cause of these changes, we analyzed fibroblast density and found that the fibroblast density is highest at the micromass periphery, where the micromass_lining cluster resides (**Fig. 1K**). Over time, micromass contraction led to a reduction in size and an increase in internal cell density, correlating with the downregulation of the myofibroblast signature (**Fig. 1K** and **L**). Spatial gene expression analysis further confirmed that *PRG4* is enriched in high-cell-density regions, whereas *THY1*, *COL1A1*, *and MFAP5* are predominantly expressed in areas of lower cell density (**Fig. 1M**). Collectively, these results suggest that increased cell density functions as a spatial signal that instructs fibroblast differentiation toward a synovial lining fate.

### Cell density controls synovial fibroblast gene expression

Next, we tested if high cell density is sufficient to induce synovial lining fibroblast differentiation. To this end, we performed bulk RNA-sequencing on synovial fibroblasts cultured at four defined cell densities: 75, 150, 300, and 600 cells/mm², respectively (**Fig. 2A**). Principal component analysis (PCA) revealed a clear transcriptional separation between low- and high-density conditions, with PC1 accounting for 80% of the variance and primarily reflecting differences in cell density (**Fig. 2B**). Differential expression analysis between high-density (300 and 600 cells/mm²) and low-density (75 and 150 cells/mm²) groups identified 899 genes with significant changes (FDR < 0.1, |log2FC| > 0.5) (**Suppl Table. 2**). Notably, high-density cultures exhibited increased expression of canonical lining fibroblast markers (*PRG4*, *PDPN*, *CLU*) and reduced expression of sublining fibroblast markers (*POSTN*, *THBS1*, *MFAP5*), with expression levels showing clear density-dependent trends (**Fig. 2C**).

**Figure 2.**
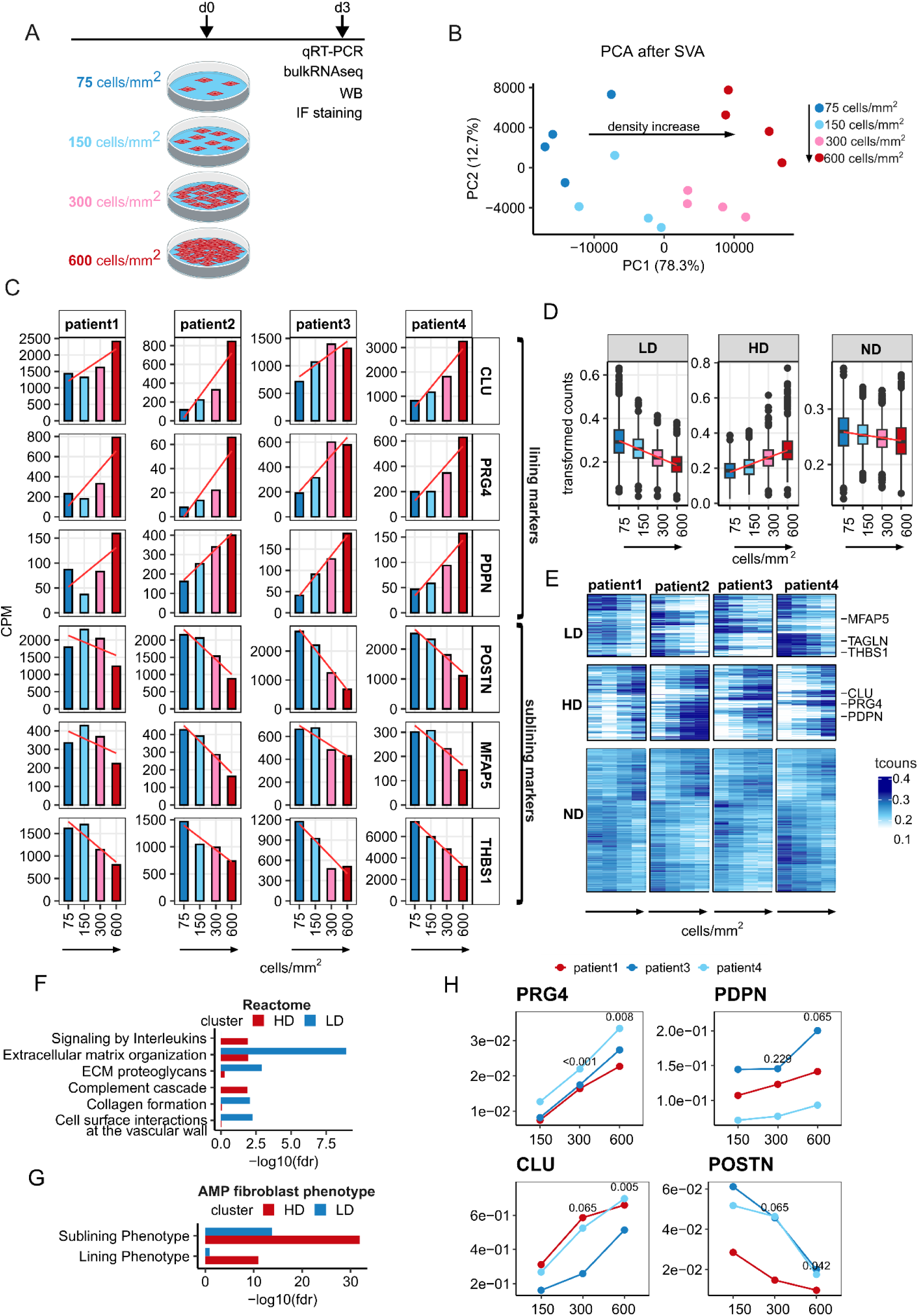
Cell density regulates fibroblast identity in cultured synovial fibroblasts. **A.** Overview of the experimental design and analysis workflow: RNA sequencing of RA patient-derived fibroblasts cultured at four different cell densities (75, 150, 300, and 600 cells/mm^2^). **B.** PCA of the normalized gene expression values after batch correction for individual cell line variability. Each point represents the expression profile of one sample. **C.** Expression changes of synovial lining (*PRG4*, *PDPN*, *CLU*) and sublining (*POSTN*, *THBS1*, *MFAP5*) markers across different cell densities. **D.** Boxplots showing gene expression patterns at four cell densities (75, 150, 300, and 600 cells/mm^2^) for three gene clusters identified by k-means clustering: low-density (LD), high-density (HD), and density-independent (ND) clusters. **E.** Heatmap showing density-dependent gene expression patterns across all samples, with genes grouped into three clusters using k-means clustering. **F.** Reactome Pathway enrichment analysis of two gene clusters identified in Fig. 2D. Colors indicate the cluster (LD/HD) used for enrichment analysis. **G.** Fisher’s exact test analysis of lining and sublining markers (defined by AMP consortium) in LD and HD clusters. **H.** qRT-PCR validation of density-dependent gene expression in synovial fibroblasts. Relative transcript levels to *GAPDH* of lining markers (*PRG4* and *CLU*) and sublining markers (*POSTN* and *CD90*) are shown. Two-way ANOVA was used to compare 300 and 600 cells/mm² with 150 cells/mm², with p-values shown above each comparison.

To identify global patterns of cell density-dependent gene signatures, we performed coseq analysis^22^ using the k-means algorithm with arcsin transformation (**Fig. 2D**, **E**, and **Suppl Table. 2**). This analysis revealed three distinct gene clusters with varying density-dependent expression patterns: a low-density gene cluster (LD) showing highest expression at 75 cells/mm^2^ with gradual decrease as density increases, containing genes such as *THBS1* and *MFAP5*; a high-density gene cluster (HD) demonstrating lowest expression at 75 cells/mm^2^ with progressive increase towards 600 cells/mm^2^, including genes such as *PRG4*, *PDPN*, and *CLU*; and a gene cluster showing no pronounced density-dependent expression pattern (ND) (**Fig. 2E**).

To investigate putative functional roles of density-dependent genes, we performed pathway enrichment analysis on these gene clusters (**Fig. 2F**). Genes induced at low cell density are enriched for pathways involved in extracellular matrix remodeling, including multiple collagens, and cell surface interaction with the vascular wall (**Fig. 2F**). In contrast, genes upregulated at high cell density were enriched for pathways associated with proteoglycan production, interleukin signaling, and the complement cascade (**Fig. 2F**). Comparison with the AMP RA/SLE Consortium single-cell atlas revealed that genes upregulated in the high-density cluster (progressively increasing from 75 to 600 cells/mm²) were specifically enriched for synovial lining fibroblast markers. Interestingly, genes associated with sublining fibroblast signatures were enriched in both low-density clusters (highest at 75 cells/mm^2^ with gradual decrease) and high-density clusters, reflecting the greater heterogeneity of sublining fibroblast populations (**Fig. 2G**). The effect of cell density on lining fibroblast marker (*PRG4. PDPN,* and *CLU*) and sublining marker (*POSTN*) expression was further validated by qRT-PCR analysis of RNA extracted from synovial fibroblasts at different densities (**Fig. 2H**). Notably, increased cell density inversely affected sublining fibroblast marker expression, supporting our observation that sublining fibroblasts are primarily localized in low-density central regions of synovial tissue (**Fig. 1F** and **2C**).

To determine whether fibroblasts can sense and respond to continuous variations in cell density, we established a density-gradient model by allowing fibroblasts to adhere to a slide tilted at a 45° angle, such that cells at the bottom experienced higher density compared to those at the top (**Fig. 3A**). We then performed spatially resolved single-cell transcriptomic analysis using *Banksy*^23^, a computational framework that integrates spatial context to refine cell type classification (**Fig. 3B–D**). Remarkably, spatially resolved single-cell analysis identified three distinct fibroblast clusters: one localized to low-density regions (L1) and two to high-density regions (H1 and H2) (**Fig. 3B–D** and **Suppl Table. 1**). The low density cluster expressed myofibroblast-associated genes, including *POSTN*, *COL1A1*, *COL5A2*, and *THY1* (**Fig. 3C** and **E**), and exhibited a proliferative phenotype marked by *TOP2A* and *MKI67* expression, whereas high-density clusters lacked proliferation marker expression (**Fig. 3C**). Notably, the H1 cluster expressed pro-inflammatory cytokines such as *CXCL12* and *CXCL6*, which were largely absent in the H2 cluster (**Fig. 3B–D**). Both high-density clusters displayed elevated expression of canonical synovial lining markers, including *PRG4*, *PDPN*, and *CLU*, which closely mirrored the transcriptional signatures observed in our bulk RNA sequencing data (**Fig. 2C**). Furthermore, correlation analysis between cell density and gene expression across the entire Xenium panel identified *CLU* and *PDPN* among the top five genes most significantly correlated with cellular density (**Fig. 3F**). We further validated these findings by applying our bulk RNA-sequencing-derived high/low density gene signatures to the spatial data (**Fig. 2** and **3G**). This analysis confirmed that the high-density gene signatures from bulk RNA-sequencing data were enriched in high-density regions of the spatial data (**Fig. 3G**).

**Figure 3.**
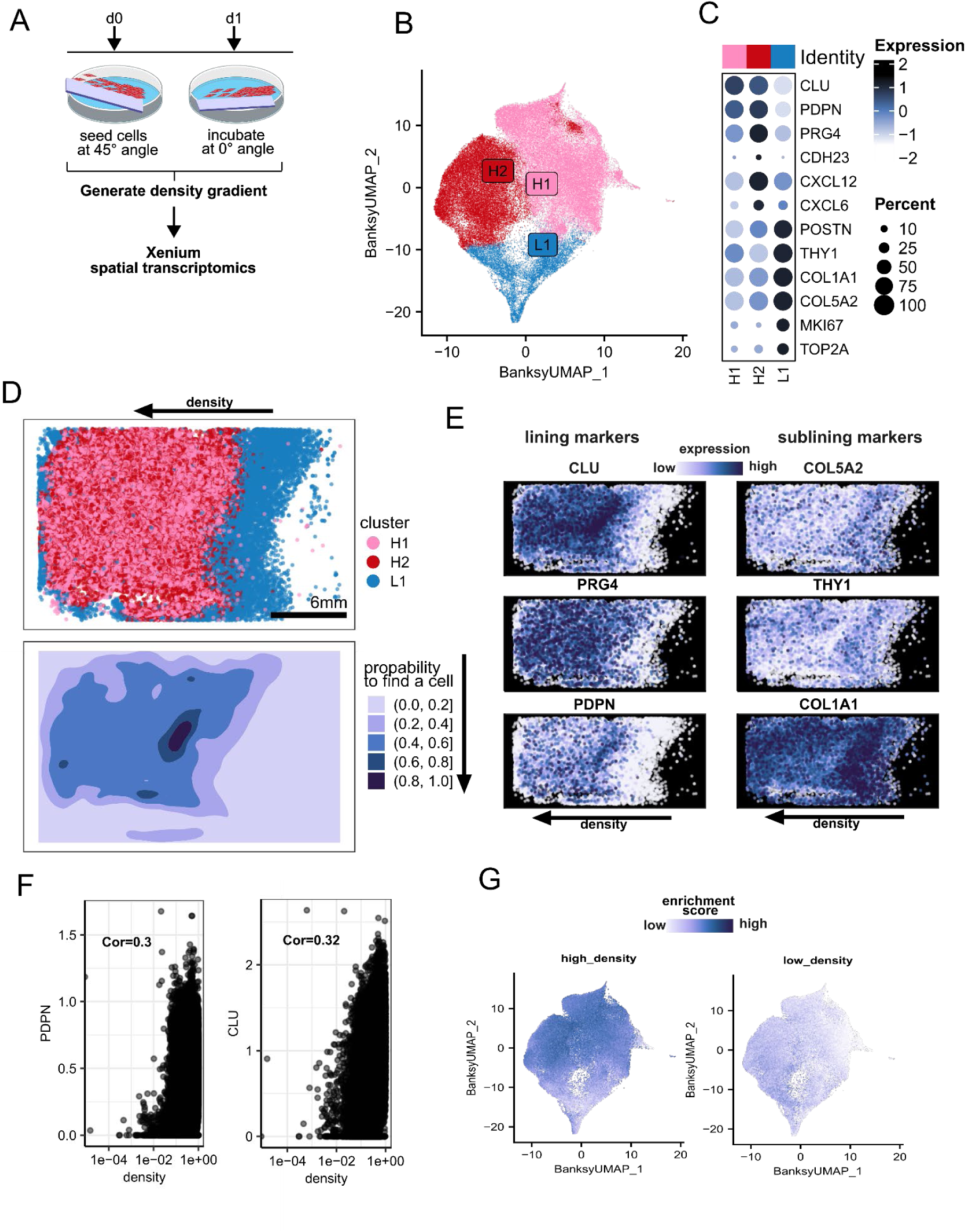
Spatial orchestration of fibroblast phenotypes through engineered 2D density gradients **A.** Experimental design of the fibroblast density gradient system. **B-C.** Spatially-resolved transcriptomic analysis of fibroblast density gradient. (**B**) Banksy UMAP visualization of fibroblast clusters identified by spatial transcriptomic analysis; (**C**) Dot plot showing key marker genes expressed across the three clusters. **D.** Spatial mapping of fibroblast clusters (H1, H2, L1) shows distinct segregation along the cell density gradient with corresponding cell density probability visualization below. **E.** Spatial visualization of marker gene expression shows lining markers (*CLU*, *PRG4*, *PDPN*) enriched in high-density regions and sublining markers (*COL5A2*, *THY1*, *COL1A1*) predominantly expressed in low-density areas, with arrows indicating direction of decreasing density. **F.** Correlation plots showing positive relationships between cell density and expression of lining markers *PDPN* and *CLU*, with each dot representing an individual cell. **G.** Banksy UMAP visualization of spatial enrichment scores for high-density (left) and low-density (right) gene signatures derived from bulk RNA-seq.

### Synovial fibroblasts sense cell density through CREB5 activation

To identify the transcriptional regulator underlying fibroblast density sensing and fibroblast quiescence, we built on our prior analysis of RA synovial tissue (**Fig. 1**), which revealed *CREB5* as a transcription factor highly enriched in high-density regions characterized by strong expression of lining fibroblast–specific genes (**Fig. 1F)**. Consistently, recent single-cell RNA-seq and ATAC-seq studies have shown elevated *CREB5* expression and motif accessibility in resting, quiescent synovial lining fibroblasts ^9,24,25^. To test if CREB5 mediates density-sensing in synovial fibroblasts, We first assessed *CREB5* gene expression and observed a consistent density-dependent induction of *CREB5* expression (**Suppl Fig. 1A**), but bulk RNA-seq showed no significant changes across the four fibroblast sublineages, likely because qRT-PCR is more sensitive (**Suppl Fig. 1B**). At the protein level, CREB5 showed modest increases (**Fig. 4A**). Since CREB family transcription factors are typically activated by phosphorylation downstream of receptor-driven kinase signaling, we next examined whether cell density influences CREB5 phosphorylation. Prior studies have identified threonine 61 (T61), a residue homologous to T71 and T53 in ATF2 and ATF7, as a potential phosphorylation site in CREB5 ^26^. With increasing cell density, we observed a marked increase in CREB5 phosphorylation at T61 (**Fig. 4A**). Notably, *CREB5* knockdown (KD) reduced pCREB5 (T61) at both densities, mirroring the reduction in total CREB5 protein, as confirmed by quantitative densitometry normalized to GAPDH (**Fig. 4A**).

**Figure 4.**
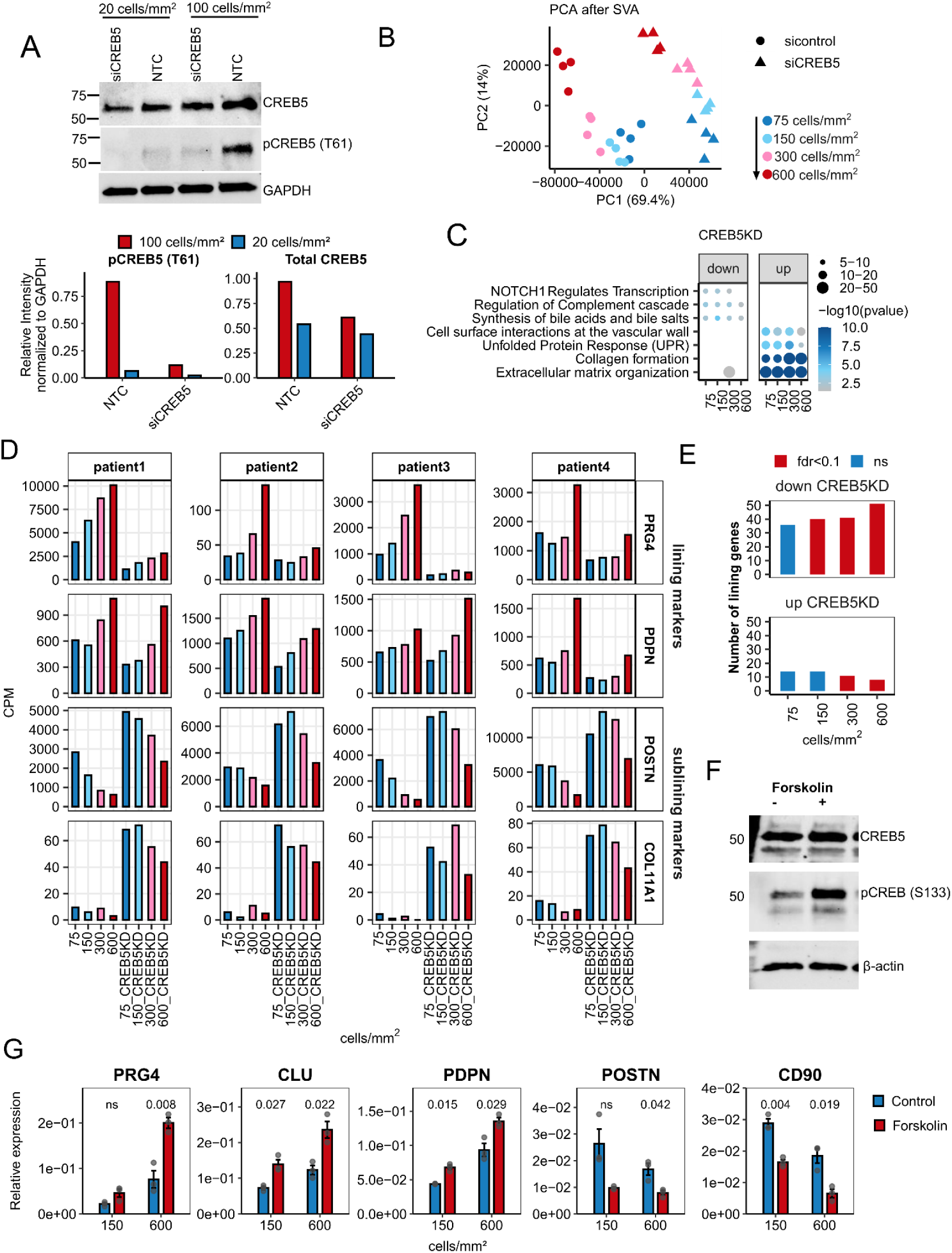
Bulk RNA-seq of *CREB5* KD in patient-derived synovial fibroblasts reveals CREB5-dependent integration of cell density in fibroblast lineage programs. **A.** Representative immunoblots of pCREB5 (T61) and total CREB5 in synovial fibroblasts after *CREB5* knockdown, cultured at low (20 cells/mm²) or high (100 cells/mm²) density, with GAPDH as loading control (top). Densitometric quantification of pCREB5 (T61) and total CREB5 normalized to GAPDH (bottom). **B.** PCA of the normalized gene expression values after batch correction for individual cell line variability. Each point represents the expression profile of one sample. **C.** GO terms enrichment analysis showing the functional pathways associated with genes upregulated or downregulated in response to *CREB5* KD across different cell densities. **D.** Expression profiles of synovial lining markers (*PRG4*, *PDPN*, *CLU*) and sublining markers (*POSTN*, *THBS1*, *COL1A1*) across four cell densities in control and *CREB5* KD conditions. **E.** Fisher’s exact test showing the enrichment of AMP-defined lining genes among diffrerntially expressed genes after CREB5 KD. **F.** Immunoblot analysis of pCREB (S133) and total CREB5 in synovial fibroblasts cultured at 100 cells/mm² for 3 days and stimulated with forskolin (10 μM, 30 min) or DMSO (0.1%, control) prior to lysis, with β-actin as loading control. **G.** qRT-PCR analysis of lining markers in Synovial fibroblast cultured at at low (20 cells/mm²) and high (100 cells/mm²) density for 6 hours followed by stimulation with forskolin (7 μM) or 0.1% DMSO control for 72 hours. Data represent biological triplicates, and *P* values are indicated above the bars.

Next, we investigated the global impact of *CREB5* KD on fibroblast density sensing by performing bulk RNA sequencing on synovial fibroblasts cultured at different densities following siRNA-mediated *CREB5* KD. PCA revealed distinct transcriptional profiles between control and *CREB5* KD , with PC1 accounting for 69.4% of the variance and primarily separating samples by KD status (**Fig. 4B**). To elucidate the functional roles of *CREB5* KD across different cell densities, we performed differential expression analysis for each condition separately (FDR < 0.1, log2FC > 1.5). Importantly, *CREB5* KD induced significant transcriptional changes at all cell densities, with 568–730 genes upregulated and 802–1139 genes downregulated, showing progressively stronger effects at higher cell densities (**Suppl Table. 2**). Although the majority of differentially expressed genes overlapped across densities, density-dependent variations in gene expression were observed (**Fig. 4B**). Pathway analysis revealed that *CREB5* KD significantly increased expression of genes involved in extracellular matrix organization, independent of cell density, particularly genes encoding collagens (*COL5A1, COL11A1, COL21A1, COL4A2*), as well as genes regulating cell cycle progression, specifically M phase regulators (**Fig. 4C**). In contrast, genes downregulated by *CREB5* KD at low density were enriched in complement cascade regulation (*C3, C1R, C2, C1*S) and metabolic pathways critical for synovial homeostasis, including chondroitin sulfate/dermatan sulfate metabolism and glycosaminoglycan metabolism, suggesting that CREB5 plays a key role in maintaining joint-protective gene expression programs (**Fig. 4C**). To identify pathway-level mechanistic insights independent of differential expression-based gene selection, we performed pathway activity analysis using PROGENy (Pathway RespOnsive GENes)^27^ with the decoupleR^28^ package to infer pathway activities from complete transcriptional signatures following *CREB5* KD (**Suppl Fig. 1C**). *CREB5* KD significantly suppressed growth factor signaling pathways, including EGFR and inflammatory pathways mediated by TNF and STAT3/JAK signaling, independent of cell density. Also, *CREB5* KD activated alternative signaling cascades including VEGF, MAPK, TGFβ, and PI3K pathways (**Suppl Fig. 1C**), suggesting that CREB5 acts as a molecular switch between inflammation and pro-fibrotic signaling in synovial fibroblasts.

Comparison of *CREB5* KD-regulated genes with synovial lining fibroblast markers from the AMP RA/SLE Consortium single-cell atlas revealed that *CREB5* KD suppressed 39-51 canonical lining fibroblast markers including *PRG4*, *PDPN*, and *CLU* across all cell densities (75–600 cells/mm²), while upregulating 25-34 sublining-associated genes including *POSTN*, *COL1A1*, and *THBS1* (**Fig. 4D** and **E**). To determine whether *CREB5* KD reverses the density-dependent lining fibroblast gene expression program, we performed unsupervised k-means clustering on genes that were differentially expressed between low (75 cells/mm²) and high density (600 cells/mm²) under control conditions (log2FC > 0.7, FDR < 0.1), and analyzed their expression patterns including high-density *CREB5* KD (600 cells/mm²). This analysis identified four distinct gene clusters (**Suppl Fig. 1E-F** and **Suppl Table. 2**): (1) a high-density-reversed cluster (HD_CREB5_rev) enriched with lining fibroblast markers, which were normally upregulated with increasing density but suppressed following *CREB5* KD; (2) a low-density-reversed cluster (LD_CREB5_rev) enriched with sublining fibroblast markers, showing the inverse density-dependent pattern that was reversed by *CREB5* KD; and (3–4) two non-changing clusters (CREB5_nc1 and CREB5_nc2) exhibiting minimal variation after KD (**Suppl Fig. 1E** and **F**). Notably, we identified EGFR, FOXO1, and SOX5 as additional regulators that were consistently downregulated across patients following *CREB5* KD, despite showing modest density-dependent induction under control conditions (**Fig. 1C** and **Suppl Fig.1D**).To our knowledge, no published studies have shown that CREB5 regulates *SOX5, FOXO1*, or *EGFR* expression. This may reflect the fact that the biology of CREB5 function remains incompletely understood. To assess the functional relevance of these transcription factors in regulating density-dependent synovial lining fibroblast gene expression *in vitro*, we performed siRNA-mediated knockdown of *FOXO1* and *SOX5* in patient-derived fibroblasts cultured at low (150 cells/mm²) and high (600 cells/mm²) densities. Suppression of either factor reduced expression of lining markers *PRG4*, *CLU,* and *PDPN* (**Suppl Fig. 1G and H**). We next extended this approach to CREB5 to directly test its role in density-dependent regulation of lining- and sublining- fibroblast gene expression. Consistent with our bulk RNA-seq analysis **(Fig. 4D**), *CREB5* KD significantly reduced expression of lining fibroblast markers (*PRG4, CLU, PDPN*) while enhancing sublining fibroblast markers (*POSTN, COL1A1, THY1/CD90*) (**Suppl Fig. 1I**). We next asked whether activation of CREB5 is sufficient to induce lining fibroblast differentiation. Because no direct small-molecule activators of CREB5 have been identified, we tested forskolin, a well-characterized activator of PKA that induces Ser133 phosphorylation^29^, a canonical regulatory site within the CREB1 family. Since our immunoblot analysis showed that pSer133 increased with density and was diminished by *CREB5* KD, we used this site as a readout of CREB5 activity. Fibroblasts treated with forskolin exhibited a robust increase in pSer133 without changes in total CREB5 protein levels (**Fig. 4F**). Forskolin treatment also induced a strong transcriptional response, characterized by upregulation of lining markers (*PRG4*, *CLU*, *PDPN*) and repression of sublining markers (*POSTN*, *THY1*/*CD90*) (**Fig. 4G**). Together, our data suggest that CREB5 activation functions as a molecular switch, promoting the lining fibroblast programs while suppressing sublining fibroblast programs.

### EGFR signaling activates CREB5 in synovial fibroblasts

We next sought to define the upstream regulation of CREB5 activation in synovial fibroblasts. To do this, we performed a targeted spatial transcriptomic screen in fibroblasts transfected with siRNAs against receptors associated with CREB activation and fibroblast function, with particular attention to EGFR, IGF1R, TRKs, and GPCRs ^30–32^ as well as Cadherin-11 (CDH11), cadherin (CDH2), and cadherin 23 (CDH23) ^33,34,35^. For each perturbation, we compared global transcriptional programs at low and high density relative to non-targeting control (siControl) to determine how knockdown of each target gene altered the density-dependent transcriptional shift.

Synovial fibroblasts were treated with siRNAs targeting 27 genes or non-targeting control. (**Suppl Table. 3**) and cultured for 6 days at low or high density with re-transfection on day 3 to ensure sustained knockdown (**Fig. 5A**). The effect of each target on fibroblast gene expression was analyzed using spatial single-cell transcriptomics with a custom probe panel optimized for measuring lining and sublining fibroblast markers. Single-cell analysis revealed that knockdown of most genes preserved the density-dependent clustering pattern observed in control siRNA-treated fibroblasts (**Fig. 5B**). However, *IGF1R* and *EGFR* KD displayed distinct fibroblast clustering patterns: *IGF1R* KD fibroblasts clustered separately at both low and high densities, whereas *EGFR* KD at high density clustered with low-density control siRNA-treated fibroblasts (**Fig. 5B** and **C**), suggesting that *EGFR* KD reverses the density-dependent transcriptional program. To quantify the magnitude of transcriptional perturbation induced by each knockdown at high density, we applied Mixscale analysis^36^, which calculates the transcriptional distance between high-density knockdown conditions and high-density control cells carrying non-targeting siRNA. Mixscale generates perturbation scores that reflect the strength of transcriptional changes induced by each knockdown relative to control. This analysis identified *EGFR* as inducing the strongest perturbation of the density-dependent transcriptional signature (**Suppl Fig. 2A**). To validate these findings, we derived high-density and low-density transcriptional signatures from control siRNA-treated fibroblasts by identifying genes significantly upregulated at high versus low density (FindMarkers, log2FC > 0.7, min.pct = 0.1). UCell algorithm scored each cell’s overall expression of these density-associated marker sets across all knockdown conditions (**Fig. 5D** and **Suppl Fig. 2B**). Notably, *EGFR* KD at high density showed a dramatic shift in the UCell signature, with high-density *EGFR* knockdown cells acquiring low-density marker scores similar to control low-density cells (**Fig. 5D**). Specifically, lining markers *PDPN* and *CLU* were significantly downregulated in high-density *EGFR* KD fibroblasts (**Fig. 5E**), confirming that *EGFR* is essential for maintaining the high-density lining fibroblast transcriptional phenotype.

**Figure 5.**
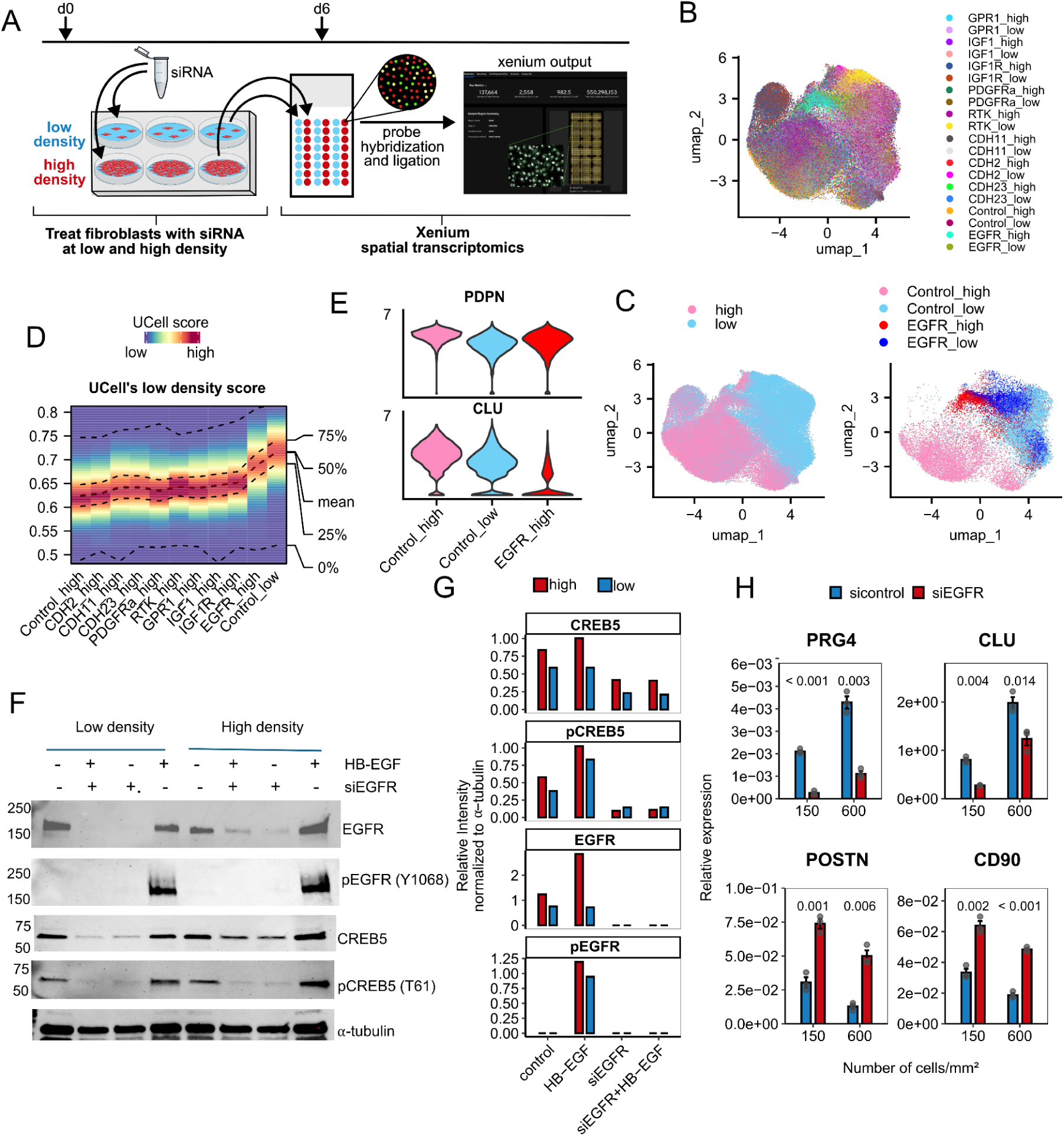
EGFR signaling regulates CREB5 activation and synovial fibroblast lineage identity in a cell density–dependent manner. **A.** Schematic diagram of the experimental design and timelines. **B.** UMAP representation of single-cell spatial transcriptomic profiles from synovial fibroblasts colored by condition (control vs. siRNA knockdown). **C.** UMAP plot of synovial fibroblasts grouped by density and *EGFR* KD condition. (Left) Cells colored by density state (high vs. low) highlight the separation of lining-like and sublining-like populations. (Right) Cells colored by experimental conditions showing control (high and low density) versus *EGFR* knockdown (*EGFR*_high and *EGFR*_low). **D.** Heatmap showing UCell scores for low-density gene signatures across all knockdown conditions at high cell density (600 cells/mm²). Each column represents a different knockdown condition or control. The color gradient indicates the strength of a low-density UCell score. **E.** Violin plots showing the distribution of *CLU* and *PDPN* expression in synovial fibroblasts across control and *EGFR* KD conditions. The x-axis represents experimental conditions and the y-axis represents normalized gene expression. **F.** Representative immunoblots of total EGFR, pEGFR (Y1068), pCREB5 (T61), and total CREB5 in synovial fibroblasts following HB-EGF stimulation or *EGFR* KD, cultured at low (20 cells/mm²) or high (100 cells/mm²) density, with α-tubulin as loading control. **G.** Quantification of EGFR, pEGFR (Y1068), pCREB5 (T61), and total CREB5 immunoblots, normalized to α-tubulin as a loading control. **H.** qRT-PCR analysis of lining (*PRG4* and *CLU*) and sublining marker (*CD90* and *POSTN*) expression in synovial fibroblasts cultured at varying cell densities under control or *EGFR* KD conditions. Data represent biological triplicates, with *P* values indicated above the bars.

Because EGFR is a surface receptor expressed on fibroblasts, we hypothesized that it may enable fibroblasts to sense their proximity to neighboring fibroblasts, a process we term *fibroblast proximity-sensing*. Consistent with this hypothesis, immunoblot and qRT–PCR analyses revealed marked upregulation of *EGFR* transcript and protein levels in fibroblasts cultured at high cell density (**Fig. 5F** and **Suppl Fig. 2C**). EGFR is known to phosphorylate and activate CREB through MAPK/ERK and RSK2 signaling ^26,37^. We therefore asked whether EGFR is responsible for CREB5 activation in densely cultured synovial fibroblasts. To test this, synovial fibroblasts were transfected with siEGFR, and CREB5 phosphorylation was assessed under low- and high-density conditions. Immunoblotting confirmed that elevated EGFR abundance in high-density cultures correlated with enhanced phosphorylation of CREB5 at T61 (**Fig. 5F and G**). Strikingly, *EGFR* KD markedly reduced pCREB5 and moderately decreased total CREB5 protein, establishing EGFR as a potential upstream regulator of density-dependent CREB5 phosphorylation (**Fig. 5F and G**). Moreover, at a transcriptional level, even a modest 1.5-to 2-fold reduction in *EGFR* was sufficient to significantly decrease the expression of lining markers *PRG4* and *CLU*, while concomitantly upregulating sublining markers *POSTN* and *THY1*/*CD90* (**Fig. 5H** and **Suppl Fig. 2C**). Together, these findings further support a role for EGFR as a fibroblast proximity sensor via activation of CREB5.

### HB-EGF induces quiescent fibroblast differentiation via the EGFR–CREB5 axis

Our model predicts that fibroblasts sense their proximity to neighboring cells through EGFR signaling. To identify potential ligands mediating this fibroblast proximity sensing, we tested the well-characterized EGFR ligands EGF, HB-EGF, and TGF-α ^38^ for their ability to induce quiescent lining fibroblast differentiation *in vitro* by upregulating lining fibroblast genes. Because CREB5 is a key downstream effector of EGFR, we first tested whether these ligands induce CREB5 activation. Immunoblot analysis revealed that HB-EGF stimulation robustly induced CREB5 phosphorylation (T61), while EGF and TGF-α stimulation produced no detectable changes compared with control (**Fig. 6A**). Quantification normalized to GAPDH confirmed the selective activity of HB-EGF (**Fig. 6B**). We next examined ligand-induced synovial fibroblast lining and sublining maker gene expression. qRT–PCR analysis showed that both EGF and HB-EGF stimulations markedly upregulated the lining fibroblast markers *PRG4* and *CLU* and reduced the sublining fibroblast markers *POSTN* and *COL1A1,* with the strongest effects observed at high cell density (**Fig. 6C and Suppl Fig. 2D**). In contrast, TGF-α showed variable responses (**Suppl Fig. 2D**). Notably, although EGF strongly induced lining-marker expression, it failed to sustain CREB5 phosphorylation (T61) (**Fig. 6A**), indicating that HB-EGF is the more potent ligand linking EGFR activation to CREB5-dependent proximity sensing. As an internal control, we assessed the expression of the canonical EGFR targets JUN and FOS, which showed robust induction following EGF and HB-EGF stimulation (**Suppl Fig. 2E**). HB-EGF stimulation also promoted phosphorylation of EGFR at Y1068, a key activation site induced by ligand engagement ^39^, accompanied by a parallel increase in pCREB5 (T61) (**Fig. 5F**). These results suggest that HB-EGF–EGFR signaling promotes CREB5 phosphorylation, coupling receptor activation to fibroblast proximity sensing.

**Figure 6.**
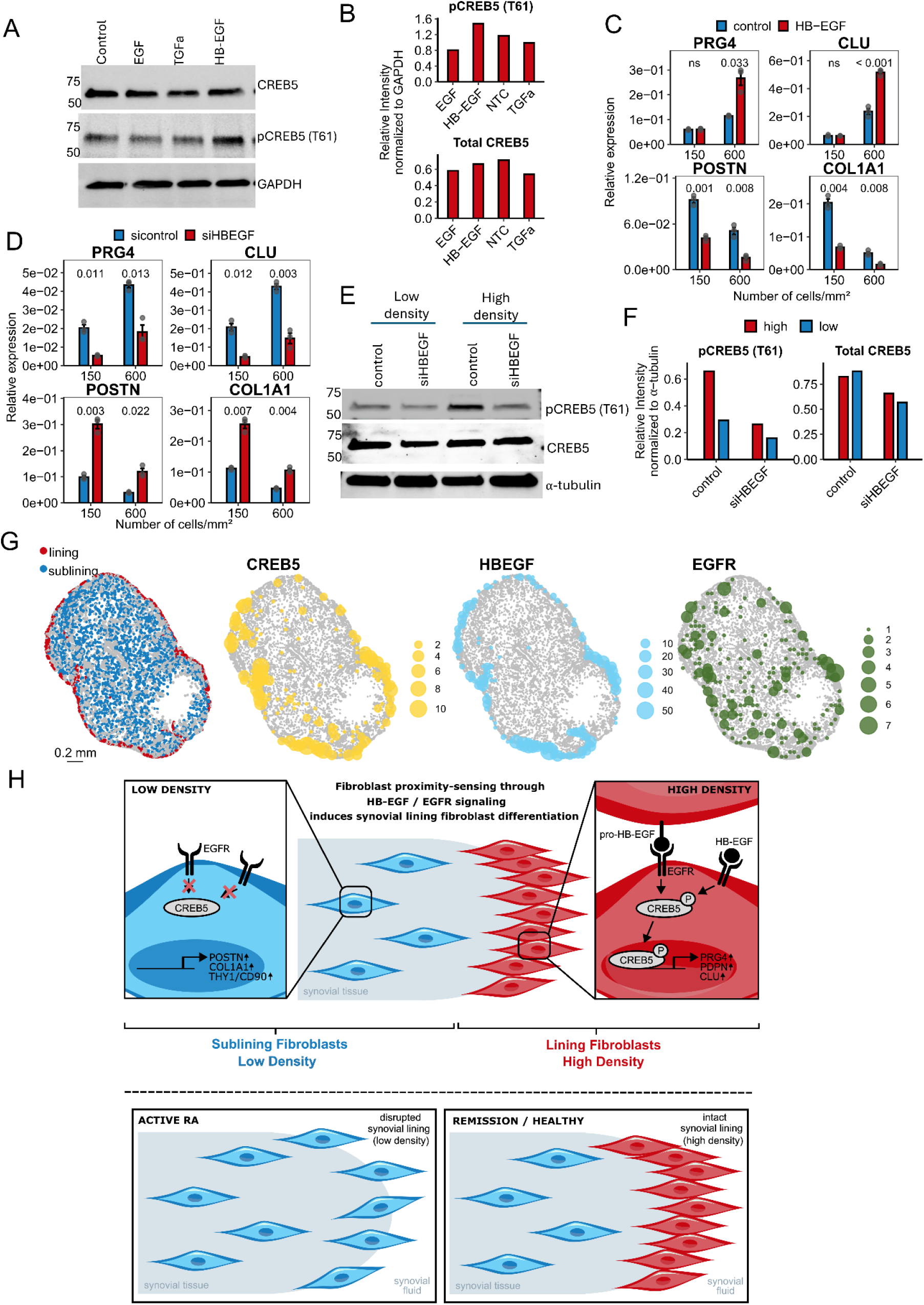
HB-EGF–EGFR–CREB5 signaling drives density-dependent lining fibroblast differentiation. **A.** Representative immunoblots of pCREB5 (T61) and total CREB5 in synovial fibroblasts cultured at high density (100 cells/mm²) following stimulation with EGF, HB-EGF, or TGFα (100 ng/ml, 10 min) or no stimulation (control), with GAPDH as loading control. **B.** Quantification of pCREB5 (T61) and total CREB5 in immunoblots of ligand stimulation, normalized to GAPDH as a loading control. **C.** qRT-PCR analysis of lining (*PRG4* and *CLU*) and sublining marker (*POSTN* and *COL1A1*) expression in synovial fibroblasts cultured at low (20 cells/mm²) and high (100 cells/mm²) densities after stimulation for 72 hours with HB-EGF (100 ng/ml), or without stimulation (control). Data represent biological triplicates, with *P* values indicated above the bars. **D.** qRT-PCR analysis of lining (*PRG4* and *CLU*) and sublining marker (*POSTN* and *COL1A1*) expression in synovial fibroblasts cultured at low (20 cells/mm²) and high (100 cells/mm²) densities after *HBEGF* KD. Data represent biological triplicates, with *P* values indicated above the bars. **E.** Representative immunoblots of pCREB5 (T61) and total CREB5 in synovial fibroblasts cultured at low density (20 cells/mm²) or high density (100 cells/mm²) following *HBEGF* KD, **F.** Quantification of pCREB5 (T61) and total CREB5 in immunoblots of *HBEGF* KD, normalized to α-tubulin as a loading control. **G.** Spatial distribution of lining and sublining fibroblasts in RA patient tissue, with gene expression mapped for *CREB5*, *HBEGF*, and *EGFR*. **H.** Model of density-dependent fibroblast differentiation through HB-EGF/EGFR-CREB5 signaling

The effect of HB-EGF on induction of synovial lining fibroblast differentiation was further examined by *HBEGF* knockdown. Consistent with *EGFR* and *CREB5* depletion, *HBEGF* KD reduced the lining fibroblast markers *PRG4* and *CLU* and decreased *EGFR* expression, while inducing the sublining markers *POSTN* and *COL1A1*, without altering *CREB5* levels (**Fig. 6D** and **Suppl Fig. 2F** ). Immunoblot analysis of synovial fibroblasts transfected with *siHBEGF* or *siControl* revealed that *HBEG*F KD markedly reduced pCREB5 (T61) levels, with a modest decrease under high cell density conditions. (**Fig. 6E**). Corresponding densitometry normalized to α-tubulin is presented (**Fig. 6F**).

Finally, to determine whether this regulatory axis is reflected in the spatial organization of the synovium during RA remission, we performed density-plot analyses comparing *HBEGF*, *CREB5*, and *EGFR* with canonical lining and sublining marker genes from 10 RA remission biopsies (**Fig. 6G**). *HBEGF* displayed strong spatial overlap with *CREB5* and lining marker genes, whereas *EGFR* exhibited a broader distribution but remained enriched within the lining region (**Fig. 6G**). These spatial relationships closely mirror our observation *in vitro*, supporting HB-EGF–EGFR signaling as a central axis driving CREB5 activity and establishing a quiescent synovial lining fibroblast identity during RA remission.

## Discussion

While RA treatment has advanced significantly over the past decades^1,2^, few RA patients are able to achieve sustained, drug-free remission. Fibroblasts play a crucial role in shaping the outcome of inflammatory diseases^20^. In RA, expansion of inflammatory fibroblasts in the sublining is a key initiator and driver of chronic tissue inflammation ^5,7,8^. Importantly, pathogenic fibroblasts differentiation in RA can directly contribute to treatment outcome, as fibrogenic differentiation of sublinig fibroblasts has been linked to treatment failure in refractory RA^10,19^. Here, we identify a novel mechanism by which synovial fibroblasts sense changes in their tissue environment to rebuild a quiescent synovial lining during RA remission. Specifically, we show that fibroblasts use cell-density cues to direct their differentiation state, providing a mechanistic explanation for synovial fibroblast heterogeneity. Our findings extend the concept of fibroblast positional identity by demonstrating that fibroblasts actively integrate signals from their surroundings, including cell–cell proximity and the availability of soluble and membrane-bound factors, to define their transcriptional and functional identity. Consistent with this model, analyses of RA remission biopsies, 2D monolayers, and 3D micromass cultures reveal that proximity sensing is a central mechanism driving lining fibroblast differentiation and restoration of a quiescent lining phenotype during remission.

Our study identifies CREB5 as the master transcriptional effector of fibroblast proximity-sensing, serving as the central node that links cell density to synovial lining fibroblast differentiation. As fibroblasts become more crowded, CREB5 mRNA and protein levels increase in proportion to spatial confinement, indicating that fibroblasts translate physical interactions within their niche into stable transcriptional changes. In this framework, CREB5 functions as the central integrator that connects tissue architecture to the synovial lining fibroblast-associated gene program. These findings indicate that CREB5 is essential for synovial lining fibroblast differentiation. Interestingly, CREB5 was recently identified as a direct transcriptional regulator of *Prg4* during establishing and maintaining the articular surface program^26^. CREB5 was specifically enriched in superficial-zone chondrocytes and was required for both TGF-β and EGFR signals to induce *Prg4* expression^26^. Members of the CREB family are broadly regulated by phosphorylation, and CREB1 in particular is a well-characterized target of mitogenic and stress-responsive pathways activated through RAS–RAF–MEK–ERK/RSK and PI3K–AKT cascades that phosphorylate Ser133 to recruit the co-activators CBP and p300 ^40,41^. However, the mechanisms controlling CREB5 activation remain poorly understood. Our systematic receptor-level analysis identified EGFR signaling as the dominant pathway coupling cell density to CREB5 activation and lining fibroblast differentiation. *EGFR* knockdown in fibroblast abolished CREB5 phosphorylation and redirected fibroblasts toward sublining fibroblast fate. These findings support EGFR as the principal upstream regulator of the density-responsive CREB5 circuit.

Among EGF-family ligands, HB-EGF emerged as the most potent activator of this pathway in synovial fibroblasts. Mechanistically, HB-EGF is unique among EGFR ligands in its ability to signal through both juxtacrine and paracrine modes. It is synthesized as a membrane-anchored pro-HB-EGF^42^ that can engage EGFR on adjacent cells via direct contact, or be cleaved by ADAM metalloproteases (ADAM10/17) to release a soluble form that diffuses locally^43–45^. At high cell density, close fibroblast–fibroblast contact increases the effective presentation of membrane-anchored pro-HB-EGF, likely enabling juxtacrine activation in which HB-EGF on one fibroblast directly engages EGFR on adjacent fibroblast. In contrast, at low density, sparse fibroblast-fibroblast contact restricts juxtacrine communication and limits the accumulation of soluble HB-EGF, leaving only weak autocrine signaling and reduced EGFR-CREB5 activation. This decrease in signal strength lowers CREB5-dependent transcription and promotes a sublining fibroblast phenotype, demonstrating that fibroblast spatial proximity directly governs ligand availability, EGFR signaling, and cell-state identity (**Fig. 6H**). Our finding is supported by a previous report demonstrating HB-EGF⁺ fibroblasts are assoicated with RA remission^14^. Our results provide the mechanistic basis for these observations by demonstrating that HB-EGF acts as a spatially restricted ligand that activates EGFR–CREB5 signaling in crowded niches to promote differentiation of quiescent lining fibroblasts. The increased EGFR and CREB5 activation observed at high density may also reflect post-transcriptional stabilization or decreased receptor turnover, consistent with the notion that spatial proximity reinforces both signal amplitude and transcriptional persistence.

Taken together, our findings reveal that synovial fibroblasts sense spatial proximity through an HB-EGF–EGFR–CREB5 axis, integrating spatial awareness, receptor activation, and transcriptional control to restore the quiescient synovial lining layer in remission RA. This model highlights the joint synovium as a self-organizing, spatially responsive tissue, in which architecture itself functions as a regulatory input, transforming cell proximity into a sustained transcriptional identity.

## Methods

### Synovial Tissue Collection and Storage

Synovial tissue samples were obtained arthroscopically from patients with early (<12 months) RA, pre- and post-treatment (with csDMARDs or adalimumab), or who were undergoing joint replacement surgery or synovectomy. Participant recruitment and tissue collection were conducted at Flinders Medical Centre (Adelaide, Australia; IRB 396.10) and Brigham and Women’s Hospital in accordance with protocols approved by the Mass General Brigham Human Research Committee (IRB 2019P002924). Excised synovial tissues were immediately cryopreserved in CryoStor® CS10 cryopreservation medium (Millipore Sigma, Cat. No. C2874), initially stored at −80°C, and subsequently transferred to liquid nitrogen for long-term storage.

### Fibroblast Cell Culture

Synovial fibroblasts were isolated from synovial tissue using a modified version of a previously described protocol ^8^. Briefly, frozen synovial tissue was thawed and rinsed in PBS, minced into small fragments, and transferred to 50 mL Falcon tubes under sterile conditions. Tissue fragments were enzymatically digested in a solution containing Dispase II (Sigma-Aldrich, Cat. No. 494207801), DNase I (Sigma-Aldrich, Cat. No. 10104159001), and Liberase TL (Sigma-Aldrich, Cat. No. 5401020001), each at a final concentration of 100 µg/mL in plain RPMI 1640 medium. Digestion was performed for 1 hour at 37°C with constant agitation on an orbital shaker.

Following digestion, 30 mL of complete FLS culture medium (Dulbecco’s Modified Eagle Medium (DMEM) supplemented with 10% FBS, 10 mM HEPES, MEM nonessential amino acids, MEM essential amino acids, 2 mM L-glutamine, 100 U/mL penicillin, 100 µg/mL streptomycin, 50 µM 2-mercaptoethanol, and 50 µg/mL gentamicin) was added. The cell suspension was filtered through a 70 µm cell strainer and centrifuged at 1500 rpm for 10 minutes. Cell pellets were resuspended in fresh complete fibroblast medium and cultured under standard conditions (37°C, 5% CO₂). The culture medium was replaced every 3 days.

For cell density experiments, synovial fibroblasts were seeded at densities of 75, 150, 300, or 600 cells/mm² in 96-well plates for quantitative reverse transcription PCR (qRT-PCR) analyses, or at densities of 20 and 100 cells/mm² in 6-well plates for Western blot assays. All experiments were performed using primary synovial fibroblasts from 2–4 independent patient donors, and each experiment was independently repeated 2–3 times across these donor-derived cell lines.

All experiments were performed using primary synovial fibroblasts of 2–4 independent patient donors, and each experiment was independently repeated 2–3 times across these donor-derived cell lines. All cells were used between passages 2-4.

### Spatial transcriptomic analysis of synovial tissue

Seurat objects containing cell type annotations and spatial coordinates were obtained from a previously published Xenium spatial transcriptomic dataset of RA synovial tissue generated in our laboratory ^8^. Objects were filtered to include only samples from RA patients in clinical remission (DAS28 < 2.6). The fibroblast Seurat object was pre-filtered to remove missegmented cells expressing myeloid and T cell markers. Standard Seurat v5 preprocessing pipeline ^46^ was applied with batch correction. Data were normalized (scale factor: 5000), variable features identified (n = 1000) and scaled prior to PCA (20 components). Harmony algorithm ^47^ was applied to correct for batch effects across patient samples (dims.use: 1:20, lambda: 1). Integrated dimensionality reduction was performed using the Harmony-corrected PCA space for k-nearest neighbor graph construction and UMAP visualization (dims: 1:20). Cells were clustered at resolution 0.5 using the Louvain algorithm. Fibroblast subtypes were annotated based on canonical marker expression: PRG4+ lining fibroblasts and THY1+ sublining fibroblasts. Additional marker genes distinguishing lining and sublining subtypes were identified using presto R package with thresholds (AUC > 0.6, adjusted p-value < 0.05, percent expressing in cluster > 0.1). Local fibroblast density was computed using kernel density estimation (KDE) with Diggle’s bandwidth selection and edge correction applied to fibroblast spatial coordinates. Density values were calculated using the spatstat R package and interpolated to assign a local fibroblast density value to each cell within the tissue section. Spatial correlation analysis was performed using the InSituCor R package^48^ to assess associations between gene expression and local fibroblast density within 50-micrometer border regions.

### Synovial Micromass Organoid Experiments

Synovial fibroblast micromass cultures were established as previously described ^21^; ^49^ with minor modifications. Briefly, cultured synovial fibroblasts were resuspended at a concentration of 200,000 cells per 35 µL in Matrigel (Corning, Cat. No. 356231). A 35 µL aliquot of the cell-Matrigel suspension was dropped into individual wells pre-coated with poly(2-hydroxyethyl methacrylate) (poly-HEMA; Millipore Sigma, Cat. No. 192066) to prevent cell adhesion. Cultures were maintained in Endothelial Growth Medium-Plus (EGM-Plus; Lonza, Cat. No. CC-5035) and imaged at defined time points. After 3–5 days in culture, micromass spontaneously detached from the substrate, forming free-floating spheroids that remained suspended in the medium for the remainder of the culture period.

### Spatial Transcriptomics

Micromass was generated (as described above) and harvested after 4, 7, 14, or 21 days and in 10% formalin overnight at 4°C, then stored in 70% ethanol at 4°C before embedding in paraffin. The Xenium slides were then prepared according to the 10X protocol (CG000580 Rev D, 10X Genomics). In brief, the slides were baked at 60°C for 2 hours and then sequentially immersed in xylene, ethanol, and nuclease-free water to deparaffinize and rehydrate the tissue. Immediately after, the Xenium slides were incubated in the decrosslinking and permeabilization solution at 80°C for 30 minutes, followed by a wash with PBS-T.The Xenium slides were then processed according to the “Xenium In Situ Gene Expression” user guide (CG000582 Rev D, 10X Genomics) for the remaining slide preparation steps. The samples were hybridized with probes from Xenium Human Multi-Tissue and Cancer Panel (PN-1000626, 10X Genomics) at 50°C for 18 hours. After hybridization, the slides were washed and incubated with a ligation reaction mix, followed by another wash step and DNA amplification. Cell segmentation staining was then performed and the slides were treated with an autofluorescence quencher and DAPI before loading into the Xenium instrument. Identical single-cell preprocessing and analysis approaches as described for synovial tissue were applied for spatial analysis of micromass organoids. For pseudo-bulk analysis, raw counts from internal fibroblast clusters were collapsed by sample and timepoint, summing expression values across all cells within each group. Genes with >95% zero counts were filtered to remove lowly expressed genes. Pseudo-bulk count matrices were analyzed using DESeq2 with design formula accounting for timepoint, with significance threshold FDR < 0.1 and |log2 fold change| > 0.5.

### Xenium Analyzer Setup and Data Acquisition

Processed Xenium slides (as described above) were loaded in the Xenium Analyzer and imaged, following the guidelines in the “Xenium Analyzer User Guide (CG000584 Rev B, 10X Genomics). After scanning, the Xenium slides were removed from the Xenium Analyzer, and processed with post-run H&E staining according to the Xenium In Situ Gene Expression - Post-Xenium Analyzer H&E Staining protocol (CG000613 Rev A, 10X Genomics).

### Fibroblast Density Gradient Experiments

Xenium slides (10x Genomics) were brought to room temperature in a sterile cell culture hood and rinsed with 70% ethanol, followed by air-drying at room temperature for 20 minutes. Slides were then submerged in 1 mM MgAct, rinsed again with 70% ethanol, and coated with poly-L-lysine (Millipore Sigma, #P4707-50ML) overnight at 4°C. After rinsing with ultrapure water (ddH₂O), slides were coated with 10% Matrigel in PBS and incubated overnight at 4°C.

Synovial fibroblasts were cultured to confluency, trypsinized, and resuspended at a density of 50,000 cells in 700 µL of complete FLS medium. For cell seeding, slides were positioned at a 45-degree angle against the edge of a large cell culture dish ^50^. The cell suspension was carefully pipetted from the top to the bottom of each slide, allowing the cells to flow downward along the surface. The dish was covered and incubated overnight at 37°C.

The following day, the medium was replaced with a fresh complete FLS medium, and the slides were repositioned horizontally within the dish. Cells were cultured for an additional 3 days. After this culture period, cells were fixed with 10% formaldehyde overnight at 4°C. Slides were then transferred to 70% ethanol for storage until processing for Xenium analysis (as described above).

### Spatial transcriptomic analysis of density gradients

Standard Seurat v5 preprocessing pipeline was applied as described above for synovial tissue analysis. Following preprocessing, spatially resolved single-cell analysis was performed using Banksy^23^ with parameters k_geom=15,30 and lambda=0.2 to integrate spatial context for cell clustering (resolution 0.3). Local cell density was computed using 2D kernel density estimation (MASS R package) with interpolation for each cell.

### siRNA Knockdown

Short interfering RNAs (siRNAs) targeting specific genes, along with a non-targeting control, were purchased from Thermo Fisher Scientific (Silencer Select) and are listed in Supplementary Table S1, including HB-EGF (Thermo Fisher Silencer Select, ID#10880). Synovial fibroblasts were reverse transfected with 20 µM siRNA using RNAiMAX transfection reagent (Thermo Fisher Scientific, #13778030) following the manufacturer’s protocol with a 1:5 ratio of RNAiMAX (µL) to siRNA (µM) in Opti-MEM. The siRNA–RNAiMAX complex was incubated at room temperature for 5 minutes before being added directly to cells cultured in complete fibroblast medium supplemented with 5% FBS. After a 24-hour incubation, the medium was replaced with a complete FLS medium containing 10% FBS.

For cell density experiments, 2 × 10⁵ synovial fibroblasts were transfected with either targeting siRNA or non-targeting control siRNA as described above. Following a 3-day incubation period, cells were trypsinized and seeded in triplicate at varying densities (150, 300, and 600 cells/mm²) in 96-well plates containing complete FLS medium. After an additional 3-day incubation, RNA was extracted for qRT–PCR or bulk RNA sequencing. For western blot analysis, synovial fibroblasts were seeded either at 2 × 10⁴ cells were seeded in each well for the total of five wells of a 6-well plate (for 20 cells/mm² experiments) or at 1 × 10⁵ cells in a single well of a 6-well plate (for 100 cells/mm² experiments) and transfected with either targeting siRNA or non-targeting control siRNA as described above. After a 3-day incubation period, the medium was removed and cells were re-transfected with the same siRNA. Following a total of 6 days in culture, cells were trypsinized and lysed as described in the western blot methods section.

For stimulation assays, synovial fibroblasts were seeded in triplicate at low (150 cells/mm²) and high (600 cells/mm²) densities in 96-well plates containing fibroblast medium supplemented with 5% serum. After 6 hours of adherence, cells were stimulated with 7 µM forskolin (Thermo Fisher Scientific, #J63292.MA), 100 ng/mL recombinant human EGF (#AF-100-15), HB-EGF (#100-47), or TGF-α (#100-16A), or treated with 0.1% DMSO as a vehicle control, and cultured for an additional 72 hours prior to RNA extraction.

### Quantitative Real-time PCR (qRT-PCR)

RNA was extracted from fibroblasts using TRIzol reagent (Invitrogen, cat. # 15596026) following the manufacturer’s protocol. cDNA synthesis was performed using the QuantiTect Reverse Transcription kit (Qiagen, #205314). Quantitative PCR (qPCR) was carried out using the Brilliant III Ultra-Fast SYBR Green qPCR master mix (Agilent Technologies, #600883) on an Agilent AriaMx Real-Time PCR system. A list of primers used for qRT-PCR is provided in the **Supplemental Table. 4.**

### Bulk RNA-seq

Cultured synovial fibroblasts isolated from four independent patient donors were lysed in TCL lysis buffer (Qiagen # 1031576) with 1% 2-mercaptoethanol at a ratio of 4ul lysis buffer per 1,000 cells. About 1,750 cells were sequenced per condition using the Smart-Seq2 platform at the Broad Institute (Brockton, MA).

### Differential gene expression analysis

Transcript-level abundances from bulk RNA-seq were quantified using the quasi-mapping approach with salmon^51^ using default parameters and the Gencode v44 reference transcriptome. Transcript-level abundances were aggregated to gene-level using the tximport R package^52^. Differential gene expression analysis was carried out using DESeq2 R package^53^ with the design formula ∼0 + group + patient to account for cell density conditions and inter-patient variation. Genes were considered significantly differentially expressed if the adjusted p-value (FDR) was less than 0.1 and the absolute log2 fold change was greater than 0.7.

### Principal component analysis

Normalized counts from bulk RNA-seq were corrected for patient-specific batch effects using ComBat-seq^54^ prior to PCA. PCA was performed using the PCAtools R package with 10% variance filtering on batch-corrected counts to visualize sample clustering across different cell densities and treatment conditions.

### Functional enrichment analysis

Gene Ontology (GO) enrichment analysis was performed using the clusterProfiler R package^55^ on differentially expressed genes, with an FDR cutoff of 0.1 applied to determine statistical significance.

To identify pathway-level mechanistic insights independent of differential expression-based gene selection, pathway activity analysis was performed using PROGENy^27^ (top 500 pathway-responsive genes per pathway) with the decoupleR R package^28^. Pathway activities were inferred from complete transcriptional signatures using multivariate linear models.

Synovial fibroblast signatures distinguishing lining versus sublining identity were derived from publicly available single-cell RNA-seq data (AMP RA/SLE Consortium^11^, https://immunogenomics.io/ampra2/). Marker genes were identified by comparing lining and sublining fibroblast clusters using the Seurat^46^ FindMarkers function with a threshold of FDR < 0.05 and average log2 fold change > 0.7, generating distinct lining and sublining gene signatures for downstream comparative analysis.

### Gene expression pattern analysis

Unsupervised clustering of genes with density-dependent expression patterns was performed using the coseq R package^22^. Genes were selected based on differential expression between low (75 cells/mm²) and high density (600 cells/mm²) conditions (FDR < 0.1, log2FC > 0.7). DESeq2-normalized count data were subjected to coseq analysis with arcsin transformation and k-means clustering (K=2:5).

### Xenium spatial transcriptomic siRNA screen

For high-density conditions, 1.0×10^5^ primary synovial fibroblasts were seeded into a single well of a 6-well plate. For low-density conditions, 2×10^4^ cells were seeded per well across five wells of the same plate, ensuring sufficient total cell numbers for downstream Xenium analysis. Cells were transfected with the indicated siRNAs (Supplementary Table. 33) and cultured for three days. A second round of siRNA transfection was performed on day 3, followed by an additional 3 days of incubation. At day 6, cells were fixed and processed for Xenium spatial transcriptomic profiling ^56^.

For data preprocessing the standard Seurat v5 pipeline was applied. High-density and low-density marker genes were identified from control cells using FindMarkers (log2FC > 0.7, min.pct = 0.1), and UCell^57^ algorithm scored density-associated marker expression across all conditions. Mixscale^36^ analysis quantified transcriptional perturbation by calculating distances between knockdown and control conditions.

### Western Blot

For cell density studies, two different seeding densities were used. For high-density conditions, 1 x 10⁵ cells were plated per well in a 6-well plate. Simultaneously, for low-density conditions, 2 x 10⁴ cells were plated per well, with a total of five wells used for this condition. Cells were incubated for 3 days. Following incubation, cells were lysed, and cells from the five low-density wells were combined. Cells pelleted by centrifugation and lysed in 100 µL of RIPA buffer (Thermo Fisher Scientific, #89900) supplemented with protease and phosphatase inhibitor mini-tablets (Thermo Fisher Scientific, #A32961) and a phosphatase inhibitor cocktail (Active Motif, #37492) for phosphorylation studies. The cell lysate was briefly vortexed at high speed to dissociate aggregates, incubated on ice for 30 minutes, and then centrifuged at high speed for 20 minutes at 4°C. Protein concentration was determined using the Pierce BCA Protein Assay Kit (Thermo Fisher Scientific, #23227). Between 30-50 µg of protein was subjected to SDS-PAGE (Bio-Rad, #1610734) and transferred to a PVDF membrane (Bio-Rad, #1704273) using the BIO-Rad TRANS-BLOT TURBO transfer system (Bio-Rad, #690BR324) with a dry transfer protocol for 30 minutes according to the manufacturer’s instructions. Membranes were blocked for 15 minutes in Everyblot blocking buffer (Bio-Rad # 12010020) then incubated overnight at 4°C with primary antibodies against CREB5 (Proteintech, #14196-1-AP, 1:500 dilution), p-CREB (Cell Signaling Technology, #9198, 1:500), EGFR (Proteintech, # 66455-1-Ig),p-EGFR (Cell Signaling Technology, #3777), p-ATF2(Cell Signaling Technology, # 24329, 1:300), SOX5 (Proteintech, #13216-1-AP, 1:500), FOXO1 (Proteintech, #18592-1-AP, 1:500), EGFR (Proteintech, #66455-1-Ig), GAPDH (Thermo Fisher Scientific, #MA5-15738), ɑ-tubulin (11224-1-AP), or beta-actin (Cell Signaling Technology, #3700). Following primary antibody incubation, membranes were incubated with HRP-conjugated secondary antibodies (Thermo Fisher Scientific, anti-Rabbit #32460 or anti-Mouse #31430) for 1 hour at room temperature. Blots were developed using SuperSignal™ West Femto Maximum Sensitivity Substrate (Thermo Fisher Scientific, #34095) or SuperSignal™ West Pico PLUS Chemiluminescent Substrate (Thermo Fisher Scientific, #34577) and imaged using a Bio-Rad ChemiDoc imaging system.

For stimulation assays, synovial fibroblasts were seeded at low (20 cells/mm²) and high (100 cells/mm²) densities as described above and cultured overnight in FLS medium supplemented with 5% serum. The following day, cells were stimulated with either 10 µM forskolin (Thermo Fisher Scientific, #J63292.MA) for 30 minutes, or 100 ng/mL recombinant human EGF (#AF-100-15), HB-EGF (#100-47), or TGF-α (#100-16A) for 10 minutes, or treated with 0.1% DMSO in 5% serum, prior to lysis.

For detection of EGFR phosphorylation, cells were seeded as described above in 1% serum media overnight, and the next day were treated with 100 ng/mL HB-EGF for 5 minutes in 1% serum media, prior to lysis.

## Acknowledgment

We thank members of the Wei lab, Korsunsky lab, and Brenner lab for helpful discussions. This work is supported by a NIH R01AR085028, a Rheumatology Research Foundation Innovative Research Grant, a Brigham and Women’s Hospital Department of Medicine - Broad Institution collaborative research Award, a Brigham and Women’s Hospital Llura Gund Award for Rheumatoid Arthritis Research. K.W. is supported by a NIH-NIAMS K08AR077037, a Burroughs Wellcome Fund Career Awards for Medical Scientists. K.B. is supported by a NIH-NIAMS T32AR007530-36.

## Author Contributions

Conceptualization: S.P., K.S.A., G.C., S.K., K.W.. Experimental design and data generation: S.P., S.K., G.C., K.B., V.K., M.D.W. Xenium data generation and panel design: G.C., S.A.P. Data analysis: K.S.A, G.C. Synovial tissue acquisition and relevant correlations: M.D.W. J.K.L., M.H.J, Figure generation: S.P, K.S.A., G.C., S.K. writing original draft: S.P., K.S.A., S.K. Reviewing and editing: all. Supervision: K.W., S.K. Funding acquisition: K.W.

## Supplementary Tables

Supplementary Table 1 - Marker genes for spatial single-cell fibroblast clusters in micromass organoids and engineered 2D density gradients.

Supplementary Table 2 - Differentially expressed genes across cell densities and CREB5 knockdown conditions.

Supplementary Table 3 - List of siRNAs used for knockdown studies

Supplementary Table 4 - List of primers used in qRT–PCR studies

**Supplementary Figure 1.**
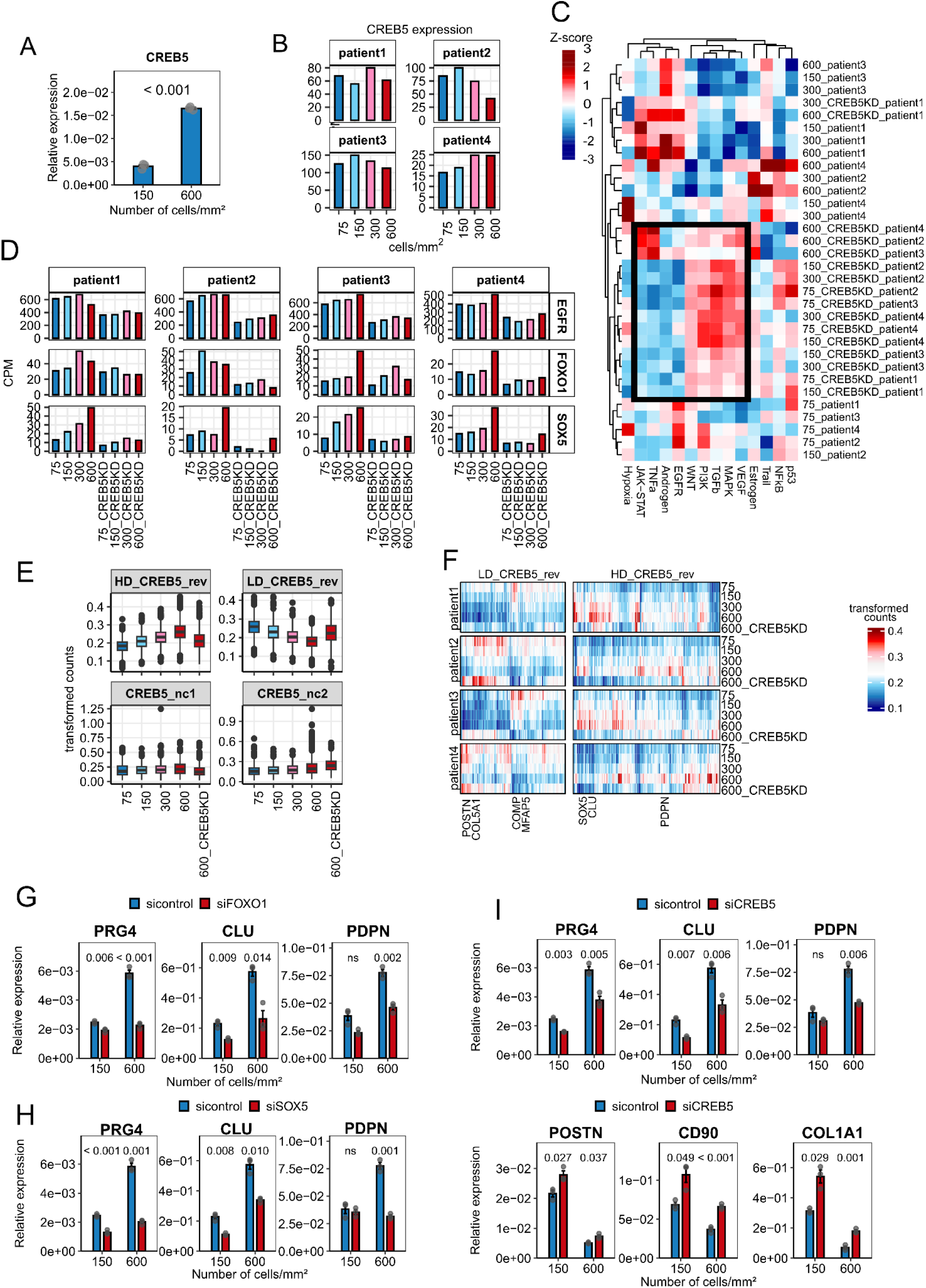
Extended bulk RNA-seq analyses of *CREB5* KD in patient-derived synovial fibroblasts confirm CREB5-dependent integration of cell density **A.** qRT-PCR analysis of *CREB5* expression in synovial fibroblasts cultured at varying cell densities. Data represent biological triplicates, with P values indicated above the bars. **B.** Expression profiles of *CREB5* across four cell densities from bulk RNA-seq (see also Fig. 2) C. Heatmap showing pathway activities across all samples and conditions using PROGENy analysis. **D.** Expression profiles of *EGFR*, *FOXO1*, and *SOX5* across four cell densities in control and *CREB5* KD conditions. **E.** Boxplots displaying normalized gene expression at three cell densities (75, 150, 300, and 600 cells/mm²) under sicontrol conditions and at 600 cells/mm² under CREB5 KD for four distinct gene clusters identified by gene pattern analysis. **F.** Heatmap showing scaled gene expression patterns across all samples for the four gene clusters identified in (E), stratified by cell density (75, 150, 300, 600 cells/mm² under sicontrol) and high-density CREB5 KD condition (600 cells/mm² under siCREB5). Each row represents a gene, columns represent individual samples. **G-H.** qRT-PCR analysis of lining marker expression (*PRG4*, *CLU*, *PDPN*) in synovial fibroblasts cultured at varying cell densities under control, *FOXO1* KD (**G**), or *SOX5* KD (**H**) conditions. Data represent biological triplicates, with *P* values indicated above the bars. **I.** qRT-PCR analysis of lining (*PRG4*, *CLU*, *PDPN*) and sublining (*POSTN*, *CD90*, *COL1A1*) marker expression in synovial fibroblasts under control or *CREB5* KD conditions. Data represent biological triplicates, with P values indicated above the bars.

**Supplementary Figure 2.**
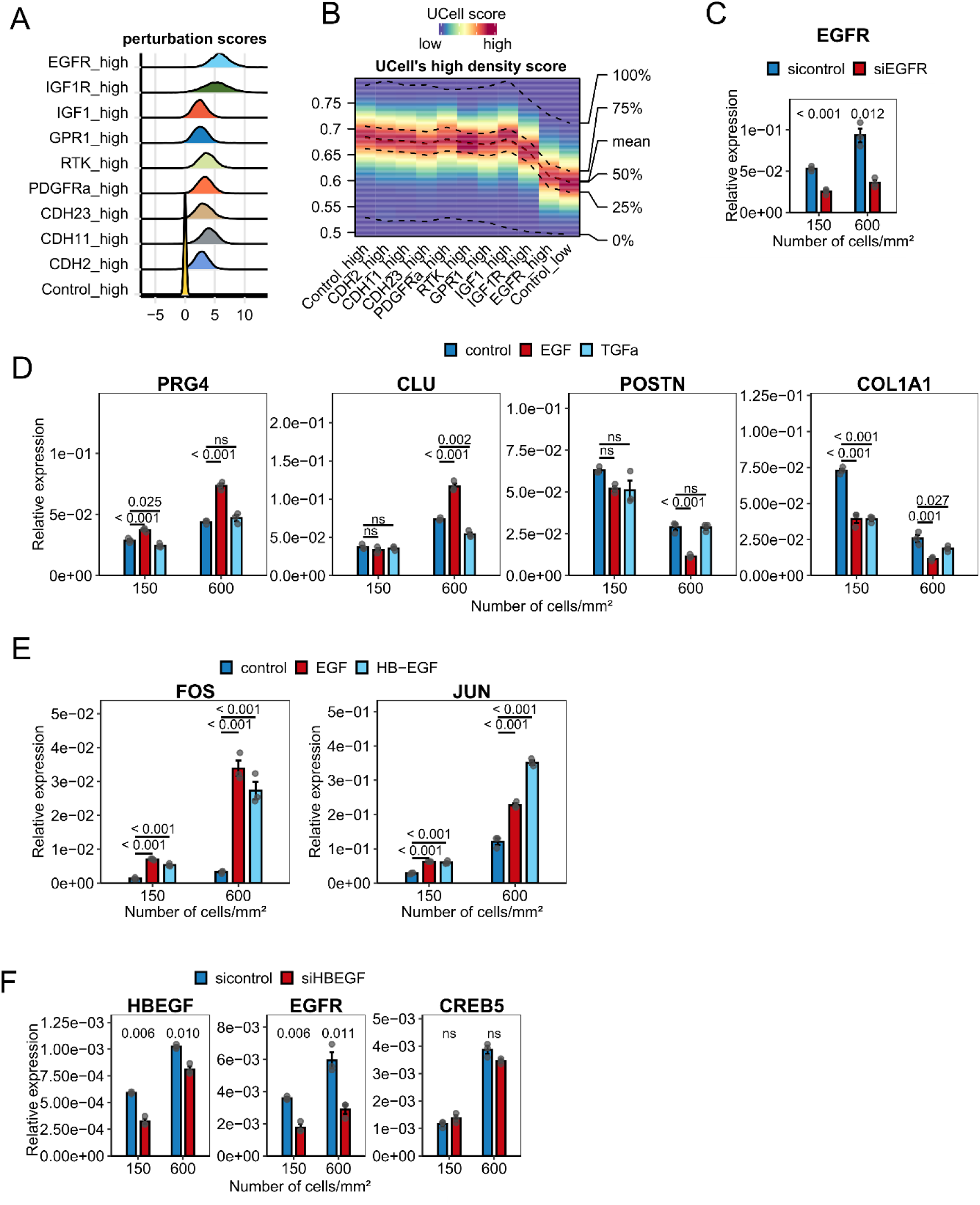
Extended analyses of the HB-EGF-EGFR–CREB5 axis in fibroblast identity. **A.** Perturbation score distributions comparing control and siRNA knockdown conditions at high cell density. **B.** Heatmap showing UCell scores for high-density gene signatures across all knockdown conditions at high cell density (600 cells/mm²). Each column represents a different knockdown condition or control. The color gradient indicates the strength of high-density UCell score. **C.** qRT-PCR analysis demonstrating the efficiency of *EGFR* KD across two cell densities, as validated by immunoblot analysis (**Fig. 5F**). **D.** qRT-PCR analysis of lining (*PRG4* and *CLU*) and sublining marker (*POSTN* and *COL1A1*) expression in synovial fibroblasts cultured at low (20 cells/mm²) and high (100 cells/mm²) densities after stimulation for 72 hours with EGF, TGFα, or without stimulation (control). Data represent biological triplicates, with *P* values indicated above the bars. **E.** qRT-PCR analysis of *JUN* and *FOS* expression in synovial fibroblasts cultured at low (20 cells/mm²) and high (100 cells/mm²) densities after stimulation for 72 hours with EGF, HB-EGF (100 ng/ml), or without stimulation (control). Data represent biological triplicates, **F.** qRT-PCR analysis of *EGFR*, *CREB5*, and *HB-EGF* expression in synovial fibroblasts cultured at low (20 cells/mm²) and high (100 cells/mm²) densities after *HBEGF* KD. Data represent biological triplicates, with *P* values indicated above the bars.

